# Condensate functionalization with motors directs their nucleation in space and allows manipulating RNA localization

**DOI:** 10.1101/2022.07.10.499452

**Authors:** Audrey Cochard, Adham Safieddine, Pauline Combe, Marie-Noëlle Benassy, Dominique Weil, Zoher Gueroui

## Abstract

The localization of RNAs in cells is critical for many cellular processes. Whereas motor-driven transport of RNP condensates plays a prominent role in RNA localization in cells, their studies remain limited by the scarcity of available tools allowing to manipulate condensates in a spatial manner. To fill this gap, we reconstitute *in cellula* a minimal RNP transport system based on bioengineered condensates which were functionalized with kinesins and dynein-like motors, allowing for their positioning at either the cell periphery or centrosomes. This targeting mostly occurs through the active transport of the condensate scaffolds, which leads to localized nucleation of phase-separated condensates. Then, programming the condensates to recruit specific mRNAs is able to shift the localization of these mRNAs towards the cell periphery or the centrosomes. Our method opens novel perspectives to examine the role of RNA localization as a driver of cellular functions.

## INTRODUCTION

The spatial organization of signaling network and biochemical reactions is of vital importance for many cellular functions. To organize the cell inner space, biomolecules and subcellular structures can be dispatched by active transport mechanisms. Long-range motor-based transport of cellular compartments along cytoskeletal networks is essential for rapid reorganization of the cellular space in response to environmental changes^1^. Microtubule-based transport is for instance necessary for the endocytic pathway, for long-distance transport of mitochondria and for lipid droplets contacts with organelles^2–4^. Although less documented than membrane-bound organelles, biomolecular condensates are also prone to interact with cytoskeletal fibers in various ways. As the main microtubule organizing center, the centrosome can be viewed as a condensate facilitating microtubule nucleation by concentrating tubulins^5^. Other examples include RNA-containing condensates such as stress granules and P-bodies, whose growth by fusion and disassembly by fission involves by microtubule-based transport^6–11^. The functional importance of condensate-microtubule interactions is also exemplified by the transport and localization of mRNAs through RNP granule transport.

Subcellular mRNA localization is a widespread process that involves mRNA transport as isolated molecules or as part of phase-separated RNP condensates^12,13^. This localization is vital for many developmental and cellular processes, from the establishment of embryo polarization to local protein synthesis at the synapses^14,15^. Motor-based positioning of specific mRNAs and subsequent local translation has for instance been described during the establishment of asymmetrical processes such as morphogen gradients in developing embryo^16–19^, cell migration^20^, neural development and synaptic plasticity^21,22^. Additionally, disruption of axonal RNP granule transport is associated with a broad range of neurodegenerative diseases^23,24^. Localizing mRNAs and RNP granules, rather than proteins, into subcellular compartments before translation favors spatially restricted protein synthesis and provides ‘outposts’ operating far from the soma^25,26^. In addition, localizing mRNAs is likely to be more energy-efficient than moving separately each protein to the right location^25^.

Due to the critical importance of RNA localization to cell fate determination, numerous methods were recently developed to describe how RNAs find their way to distinct subcellular compartments, and how this impacts RNA functions and processing. For example, the direct visualization of RNA molecules in living cells and organisms has been instrumental to elaborate our current understanding of RNA localization mechanisms^27,28^. Complementary to imaging approaches, transcriptomic RNA sequencing-based methods also described a variety of RNAs enriched in specific subcellular areas^29–31^. Motor proteins from all three families, i.e., kinesin, dynein, and myosin, have been identified as the drivers of short- and long-range mRNA transport along the cytoskeleton^32,33^. Though further studies are necessary to decipher the building blocks required to recruit, direct and release specific mRNAs to a particular destination, one recurrent scenario involves RNA binding proteins (RBP) and motor adaptors, linking mRNAs and motor proteins^12,34–37^.

Beside its role in guiding long-range transport, the cytoskeleton also contributes to the mechanical integrity of cells. Due to its inherent heterogeneity and dynamic nature, determining how such a meshwork impacts RNP condensation remains difficult to quantify. Yet, some biophysical implications of the cytoskeleton meshwork on phase separation mechanisms have recently started to be investigated both theoretically or experimentally. For example, the cytoskeleton modeled as an elastic meshwork, and acting at length scales comparable to condensate sizes, has been seen to modify nucleation and coarsening of phase separation systems^38–40^. In the very large *Xenopus* oocytes, the actin meshwork provides steric hindrance limiting nucleolar fusion as well as counter balancing sedimentation by gravity^41^. In epithelial cells, it has been shown that weak and non-specific interactions between cytoskeleton elements and the condensate surface may account for mutual influences^42^. One missing element in this description is the effect of molecular motors on phase-separated condensates to explore how transport could shape condensate formation and localization. To fill this gap, and examine how motor proteins could impact RNP phase separation, we adopted an approach allowing the reconstitution in cells of motor-functionalized condensates.

Novel tools have been developed allowing the formation of artificial condensates with programmable properties in cells. Indeed, such technologies bring novel perspectives both for addressing new biological questions and for further biotechnological improvements^43–54^. In this context, we engineered artificial condensates made of protein scaffolds that are prone to phase separate and functionalized them to interact with microtubule-bound motor proteins. Our first aim was to examine how motor proteins would affect condensate formation and localization. A second goal was to build minimal RNP condensates recruiting a unique RNA, making it possible to explore condensate-mediated RNA delocalization.

Condensates are thought to form through liquid-liquid phase separation (LLPS) induced by weakly interacting multivalent biomolecules. Based on this observation, our system relies on a self-interacting multivalent protein driving the formation of the condensates and fused to microtubule-interacting domains (from either a motor protein or a motor adaptor). We previously developed the ArtiGranule system, which relies on multivalent cores of ferritin monomers cross-linked by the self-interacting domain F36M-FKBP (Fm)^46,54^. Here, we replaced the ferritin core by a multimerization domain consisting of five consecutive Fm repeats (5Fm)^55^. We investigated two plus-end motors (KIF1A and KIF5B), one minus-end motor (KIFC1) and one adaptor of the dynein motor protein (BICD2). We first showed that the resulting scaffold proteins underwent LLPS in cells and that condensates functionalized with plus-end kinesins (thereafter called plus-end motor-condensates) were robustly positioned at the edge of cells. In contrast, condensates functionalized with the minus-end kinesin or the dynein adaptor (thereafter called minus-end motor-condensates) eventually formed a unique body at the centrosome. Interestingly, the localization of condensates was determined at the nucleation step. Our observations support a two-step process; first, motors moved quickly either towards the cell periphery or the centrosome, depending on the motor; then this led to the local accumulation of the multivalent protein on microtubules, and eventually to the formation of asymmetrically positioned large condensates through phase separation. In the case of BICD2, we additionally observed some condensate nucleation throughout the cytosol, followed by their directed transport to the vicinity of the centrosome and their coalescence.

In addition to our assay based on constitutive interactions between condensates with microtubules, we also developed a system where condensate interaction with motors or dynein adaptors could be chemically triggered using a chemically-inducible dimerization strategy^56^. Here, we observed that, upon induction of their interaction with the cytoskeleton, preformed condensates re-localized at the cell periphery or at the centrosomes, depending on the directionality of the motors.

Finally, we engineered motor condensates programmed to recruit either exogenous or endogenous mRNAs, using the MS2-MCP (MS2 Coat Protein) system. We found that bi-functionalized condensates, with both MCP and motor proteins (motor/MCP condensates), were asymmetrical positioned in cells and recruited heterologous MS2-containing mRNAs. We then studied the ASPM mRNA, which normally localizes at the centrosome during mitosis^57^ and showed that our motor/MCP condensates successfully perturb the spatial distribution of endogenous MS2-tagged ASPM mRNA.

## RESULTS

### Plus-end motor condensates localize at the periphery of cells

To build model condensates functionalized with plus-end kinesin motors in living cells, we generated a chimeric construct composed of two functional parts: a multivalent protein domain triggering LLPS in cells, fused to a kinesin motor domain to ensure trafficking along microtubule tracks. As a multivalent protein domain, we designed the 5Fm module, composed of five repetitions of the dimerizable mutant F36M of the FKBP protein (Fm) (Fig. 1A)^55^. Expression of emGFP-5Fm and mCh-5Fm in HeLa cells for 24h led to the formation of hybrid micrometric condensates composed of both emGFP and mCherry fusion proteins and randomly localized throughout the cytosolic space (Fig. 1B, left panel). For the kinesin motor domain, we first considered a truncation of the human kinesin-3 KIF1A (aa 1-383), which ensures the processivity of the motor (Fig. 1A)^58,59^. KIF1A(1-383) was fused to emGFP-5Fm (giving rise to the KIF1A-emGFP-5Fm plasmid), and the localization of the fusion protein was compared to the control LLPS scaffold emGFP-5Fm lacking any motor. Interestingly, when both the motor-LLPS scaffold (KIF1A-emGFP-5Fm) and the LLPS scaffold (mCh-5Fm) were co-expressed in HeLa cells during 24h, chimeric condensates were mostly found localized at the vicinity of the cell periphery, next to the membrane. In these conditions, almost all cells displayed highly asymmetrical localization patterns, often consisting of 3-5 micrometric condensates per cell (Fig. 1B, middle panel). To quantify the degree of asymmetry among cells, we measured the fraction of mCherry fluorescence, *i.e.* of the non-motor part of the scaffold, in the 25% peripheral area of the cells (I_25_) (Methods)^58^. Using the motor-less scaffold, as expected, condensates did not display any asymmetrical positioning and gave a I_25_ value of 20% +/- 9% (mean +/- SD, Fig. 1C). In contrast, for the chimeric condensates containing the KIF1A(1-383) motor, the I_25_ value was higher (34 +/- 9 %), in accordance with the visualization of asymmetric patterns. Therefore, KIF1A condensates are efficiently localized at the cell periphery.

**Figure 1:**
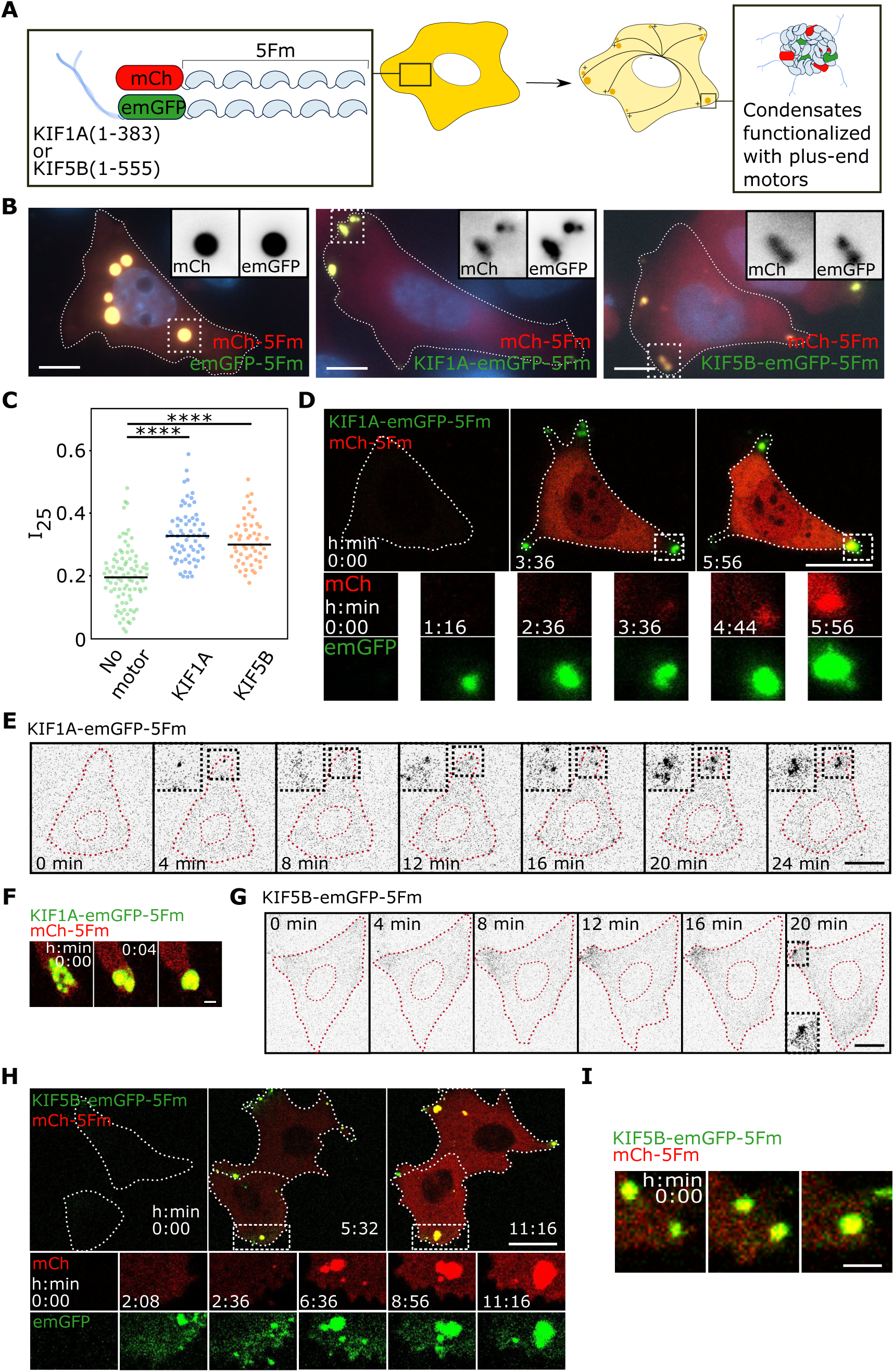
Functionalization of artificial condensates with plus-end motors drives their localization at the cell periphery. **A.** Schematic of the expected peripheral localization of condensates following transfection of mCh-5Fm and KIF1A-emGFP-5Fm or KIF5B-emGFP-5Fm (Fm = F36M-FKBP) in HeLa cells. **B.** Representative epifluorescence imaging of three cells expressing non-functionalized condensates (left panel), and KIF1A of KIF5B condensates (middle and right panel, respectively). Nuclei were stained with DAPI (blue). Grayscales inserts correspond to the red (mCh) and green (emGFP) channels of the regions delineated by dashed squares. Scale bar, 10 µm. **C.** Distribution of the fraction of mCherry fluorescence in the peripheral 25% of the cell (I_25_) for cells displaying non-functionalized condensates (left), and KIF1A or KIF5B condensates (middle and right, respectively), with each dot representing one cell (N = 87, 67 and 53, respectively). Differences between no motor and KIF1A or KIF5B were statistically significant using a Wilcoxon rank-sum test (****: p-values < 10^-11^). **D.** Time-lapse confocal imaging of the formation of KIF1A condensates in a cell (delineated by a dashed line), starting 4 h after transfection. The dashed squares indicate the region enlarged in the time-lapse images below (separate red and green channels). Scale bar, 20 µm. **E.** Epifluorescence imaging of the early time points of KIF1A-LLPS scaffold expression. Scale bar, 10 µm. **F.** Confocal imaging of coalescence events of KIF1A condensates. Scale bar, 2 µm. **G.** Epifluorescence imaging of the early time points of KIF5B-LLPS scaffold expression. Scale bar, 10 µm. **H.** Same as (D) for KIF5B condensates. **I.** Confocal imaging of coalescence events of KIF5B condensates. Scale bar, 2 µm.

In order to extend our assay, we next examined cells expressing a second plus-end directed motor domain fused to LLPS scaffolds, KIF5B(1-555)-emGFP-5Fm (Fig. 1A). The KIF5B(1-555) truncated proteins contains the motor and neck domains and part of the coiled-coil domain of mouse KIF5B^60^. As with KIF1A-LLPS scaffolds, epifluorescence imaging of HeLa cells 24 h after co-transfection of KIF5B-LLPS scaffolds (KIF5B-emGFP-5Fm) and LLPS scaffolds (mCh-5Fm) showed an asymmetrical localization of condensates at the periphery of cells (Fig. 1B, right panel). In accordance with these observations, the I_25_ value was 32% +/- 8% (mean +/- SD, Fig. 1C). Altogether our data showed that plus-end kinesin condensates are robustly positioned at the edge of cells.

To verify that the motor domain needs to be part of the LLPS scaffold for the condensate to be relocated, we examined the localization of non-functionalized LLPS scaffold (mCh-5Fm) in the presence of motor domains lacking the 5Fm multivalent domain (KIF1A-emGFP or KIF5B-emGFP). In these conditions, mCherry condensates were randomly dispersed throughout the cytosol, (Fig. S1A, red), whereas the motor domains accumulated in some regions of the cell periphery as expected (Fig. S1A, green). As a second control, we assessed the importance of multivalent LLPS scaffold interactions in the localization of plus-end motor condensates. By design, condensates should be disrupted by adding a competitive ligand of Fm dimerization (FK506). HeLa cells were transfected with KIF1A-emGFP-5Fm and mCh-5Fm, and FK506 was added at micromolar range either directly after transfection (to probe condensate formation) or 24 hours after transfection (to probe condensate dissociation). In both situations, after 26 hours of transfection, we found an absence of condensates (Fig. S1B, red) and a motor domain signal that either formed a gradient of concentration towards some regions of the cell periphery or was homogeneously diffuse in the cytoplasm (Fig. S1B, green). Time-lapse microscopy following addition of FK506 24 h after transfection showed a very fast dissolution of the condensates, with the LLPS scaffold (mCherry signal) diffusing in the whole cell in a few seconds and the motor-LLPS scaffold (emGFP signal) losing its condensed state but remaining at the cell periphery. Our method thus allows for the controlled inhibition and disassembly of plus-end motor condensates upon drug addition.

### Dynamics of formation and localization of plus-end motor condensates

While condensate positioning at the cell periphery is consistent with an active transport mediated by plus-end motors along microtubules, the localization kinetics remained to be determined. One possible chronology is, first, condensate nucleation throughout the cytosol, followed by condensate transport along microtubules. Alternatively, transport of LLPS scaffolds powered by motors could first induce their accumulation at peripherical sites, which would then trigger local condensate nucleation. To distinguish between the two scenarios, we monitored the early times of condensate formation using time-lapse microscopy (Supplementary movie 1). 4 h after co-transfection of KIF1A-LLPS and LLPS scaffolds (KIF1A-emGFP-5Fm and mCh-5Fm), we firstly noticed the strong accumulation of the motor scaffolds at the tips of the cells, illustrating the capacity of kinesins to localize within short time scales (Fig. 1D). Very often, the fast peripheral nucleation of KIF1A condensates occurred as soon as fluorescent KIF1A-LLPS scaffolds became detectable in the cytosol (Fig. 1E). These condensates tended to grow to eventually form large spherical bodies that could reach few micrometers. In addition, nearby condensates tended to coalesce (Fig. 1F).

Interestingly, we found that KIF1A-LLPS scaffolds condensed systematically ahead of LLPS scaffolds, with LLPS scaffolds then accumulating in preformed KIF1A condensates (Fig. 1D). This contrasts with the intrinsic ability of LLPS scaffolds to form randomly localized condensates, as seen in cells expressing KIF1A-emGFP lacking the LLPS domain (Fig. S1A). Altogether, these observations indicate that KIF1A condensates recruit the non-functionalized LLPS scaffolds, thus preventing their independent phase separation. In addition to peripheral condensate nucleation, we also observed rare events of long-ranged (2 to 5 µm/min) condensate transports towards the periphery (Fig. S1C).

The same dynamic characteristics were found when observing the formation of KIF5B condensates (KIF5B-emGFP-5Fm and mCherry-5Fm) (Supplementary movie 2). Early observations of KIF5B-LLPS scaffolds showed an immediate asymmetrical pattern, with a sharp gradient of fluorescence forming at the membrane and shortly preceding nucleation events (Fig. 1G). The condensation of KIF5B-LLPS scaffolds at the cell periphery also occurred ahead, followed by the recruitment of LLPS scaffolds (Fig. 1H). Condensates in close proximity tended to coalesce (Fig. 1I). As for KIF1A condensates, rare directed transport events were observed (Fig. S1D).

Taken together, our observations showed that condensate positioning at the cell periphery occurred predominantly by nucleating phase separation directly at the final sites rather than by transporting already formed condensates to their destination.

### Minus-end motor condensates localize at the centrosomes

We next examined the positioning of condensates using minus-end motors conjugated to LLPS scaffolds. We first used the human KIFC1(125-673) truncation that includes the coiled coil and motor domains required for motor processing^60^. KIFC1(125-673) was fused to emGFP-5Fm, and the resulting emGFP-5Fm-KIFC1 scaffold was co-transfected along with the LLPS scaffold (mCh-5Fm) in HeLa cells (Fig. 2A). Strikingly, after 24 h of expression, most cells displayed a single condensate in the cytosol near the nucleus (mean number of condensates per cell = 1.4 +/- 0.9, Fig. 2B, middle panel, and Fig. 2C). This contrasted with control cells transfected only with LLPS scaffolds (emGFP-5Fm and mCh-5Fm), which displayed in average 4 condensates per cell (mean +/- SD = 4.0 +/- 3.0, Fig. 2B, left panel, and Fig. 2C). As an alternative to minus-end kinesin motor, we also assessed the mouse dynein adaptor BICD2 (aa 15-595)^60^. We co-transfected BICD2-emGFP-5Fm along with the LLPS scaffold (mCh-5Fm) in HeLa cells (Fig. 2A). As observed with KIFC1, most cells displayed after 24 h a single condensate localized near the nucleus (mean number per cell = 1.2 +/- 0.6, Fig. 2B, right panel, and Fig. 2C). Interestingly, for both KIFC1 and BICD2, the single condensates docked at the centrosomes, as demonstrated by immunostaining of pericentrin (Fig. 2D). Altogether, minus-end motor functionalization of LLPS scaffold robustly led to the formation of a single condensate at the centrosome.

**Figure 2:**
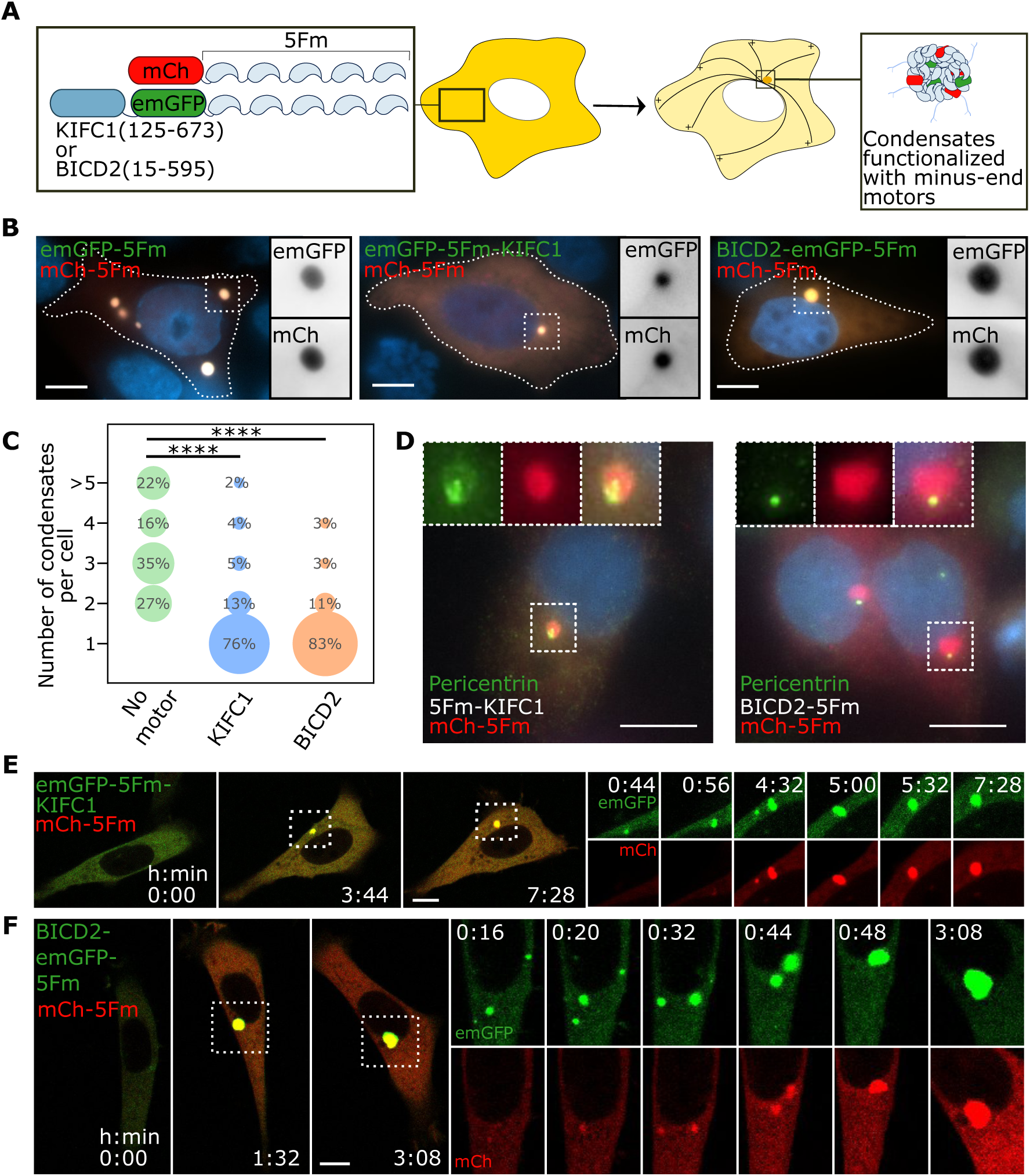
Functionalization of artificial condensates with a minus-end motor or a dynein adaptor drives their localization at the centrosome. **A.** Schematic of the expected centrosomal localization of condensates following transfection of mCh-5Fm and emGFP-5Fm-KIFC1 or BICD2-emGFP-5Fm (Fm = F36M-FKBP) in HeLa cells. **B.** Representative epifluorescence imaging of three cells expressing non-functionalized condensates (left panel), and KIFC1 or BICD2 condensates (middle and right panel, respectively). Nuclei were stained with DAPI (blue). Grayscale inserts correspond to the red (mCh) and green (emGFP) channels of the region delineated by dashed squares. Scale bar, 10 µm. **C.** Percentage of cells displaying 1, 2, 3, 4 or more than 5 condensates, for cells expressing non-functionalized condensates (left, N = 89 cells), and KIFC1 or BICD2 condensates (middle, N = 85, and right, N = 73, respectively). Differences between no motor and KIFC1 or BICD2 were statistically significant using a Pearson’s chi-squared test (****: p-values < 10^-25^). **D.** Epifluorescence imaging of cells displaying a KIFC1 or BiCD2 functionalized condensate (red, left and right, respectively) after immunostaining of pericentrin (green) as a centrosome marker. Nuclei were stained with DAPI (blue). Scale bar, 10 µm. **E.** Time-lapse confocal imaging of the formation of a KIFC1 condensate in a cell, starting 4 h after transfection. The dashed squares indicate the region enlarged on the right. Scale bar, 10 µm. **F.** Same as (E) for a BICD2 condensate. Scale bar, 10 µm.

As expected, the control co-expression of motor constructs lacking the LLPS multivalent domain (emGFP-KIFC1 or BICD2-emGFP) and non-functionalized LLPS scaffold (mCh-5Fm) led to cells displaying mCherry condensates randomly distributed though the cytosol (Fig. S2A). Of note, unlike the efficient peripheric localization of KIF1A-emGFP and KIF5B-emGFP, we observed little centrosomal accumulation of emGFP-KIFC1 or BICD2-emGFP. As for the plus-end kinesin scaffolds, adding the Fm competitor ligand FK506, immediately or 24h after transfection, suppressed condensates (Fig. S2B). These controls demonstrated the requirement of motor-LLPS scaffolds to localize condensates, as well as the need of a multivalent scaffold to trigger LLPS.

### Dynamics of formation and localization of minus-end motor condensates

To examine the pathway leading to the emergence of single condensates at the centrosomes, we monitored the early steps of their formation, starting 4 hours after transfection of emGFP-5Fm-KIFC1 or BICD2-emGFP-5Fm along with mCh-5Fm (Supplementary movies 3 and 4). In both cases, we found that condensates primarily nucleated at the vicinity of the nucleus (Fig. 2E-F). This led to the emergence of a single condensate that kept on growing, including by coalescence of smaller condensates appearing nearby. As observed with the plus-end motors, the non-functionalized scaffold accumulated exclusively at the site of motor condensates (Fig. 2E-F). In the case of BICD2 condensates, we also observed long-range transport of condensates nucleated far from the nucleus, coalescing into one large condensate during transport (Fig. 2F and S2C).

Taken together, our observations showed that minus-end motor condensates mainly nucleate at the vicinity of the nucleus, and then recruit non-functionalized scaffolds.

### The timing of non-functionalized scaffold enrichment into motor condensates depends on their localization in cells

One interesting feature shared by the four motor condensates is their ability to capture the non-functionalized LLPS scaffolds. Yet, the intracellular space being very heterogenous in term of physical properties, such as crowding and geometry, condensates’ subcellular location may impact some of their characteristics. To examine further this aspect, we studied more closely the enrichment of non-functionalized scaffolds into condensates depending on their localization in cells. We quantified the delay between the initial nucleation of motor condensates and the first discernible enrichment of the non-functionalized LLPS scaffold. We found that co-localization occurred after 1 to 2 hours using the plus-end motors (mean 83 min with a coefficient of variation CV of 40% and 74 min with a CV of 39% for KIF1A and KIF5B, respectively, Fig. 3A and 3B), contrasting with less than 20 min using the minus-end motors (mean 8 min with CV of 163% and 18 min with CV of 89% for KIFC1 and BICD2, respectively, Fig. 3A and 3C). Therefore, the delay of LLPS scaffold enrichment into preformed motor condensates was much longer for plus-end than minus-end motors. This difference in temporality may result from two non-exclusives factors: first, plus-end and minus-end motor condensates localized in two different areas of the cell where the pool of available LLPS scaffold may strongly differ because of the cell geometry, narrower at the periphery than close to the nucleus (Fig. 3D). Additionally, molecular crowding may strongly vary between the centrosome and the cell membrane area. Secondly, the processivity of our plus-end and minus-end motor differ, with only the plus-end motor scaffolds leading a rapid leap in concentration and condensate nucleation (Fig. 1E and 1G). Nucleation could thus occur before the non-functionalized LLPS scaffold reaches a sufficient concentration for enrichment. Altogether our data show that the timing of the enrichment of non-functionalized scaffolds into motor condensates depends on their localization in cells.

**Figure 3:**
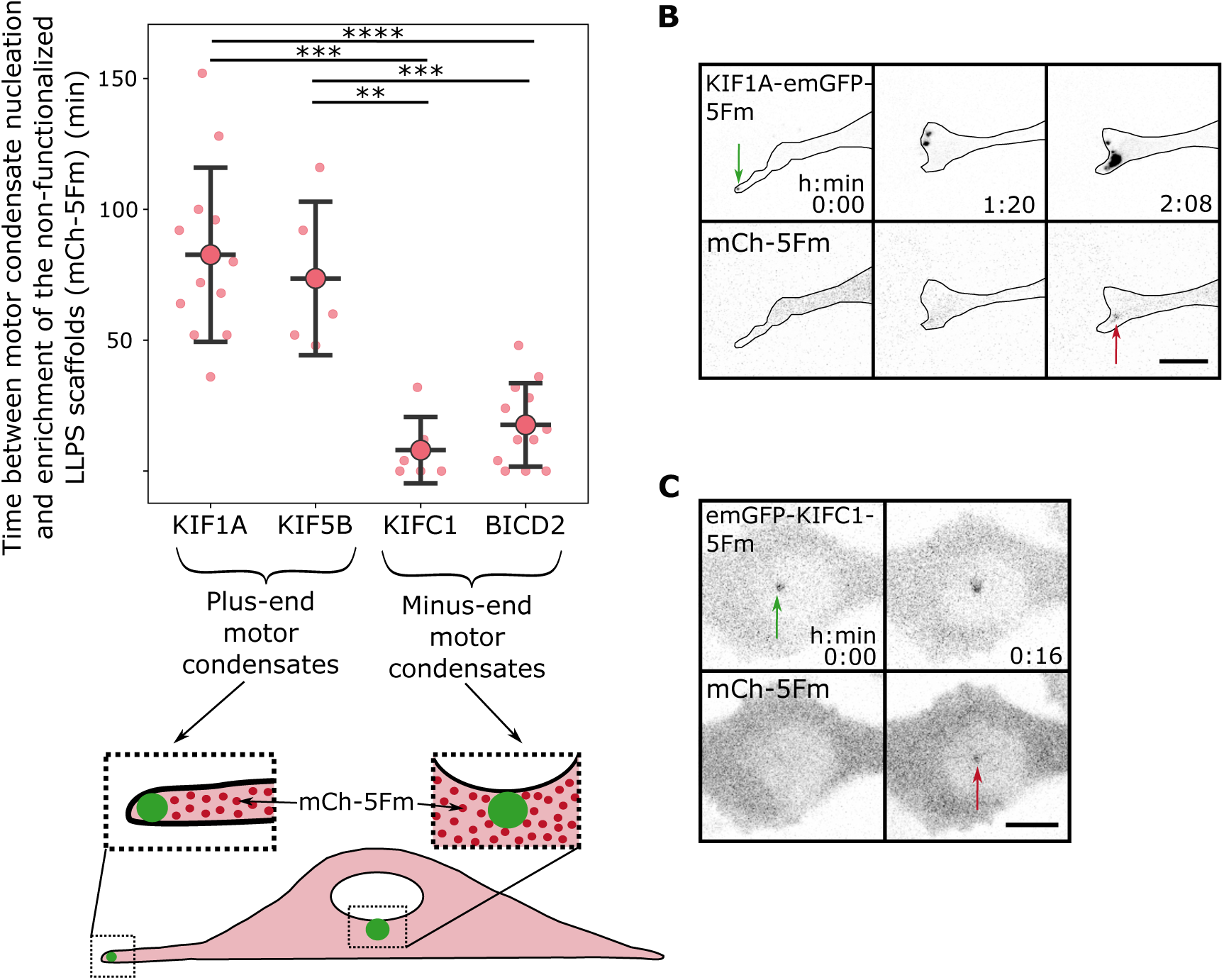
The enrichment of non-functionalized scaffolds in motor condensates differs depending on their cellular localization. **A.** Top: Delays between nucleation of the motor-LLPS scaffold and first detectable non-functionalized scaffold (mCherry signal) enrichment in condensates, for plus-end motors KIF1A (N = 12 cells) and KIF5B (N = 5), minus-end motor KIFC1 (N = 12) and dynein adaptor BICD2 (N = 6). Differences between plus-end motors and minus-end motor / motor adaptor were statistically significant using a Wilcoxon rank-sum test (**: p < 10^-2^; ***: p < 10^-3^; ****: p < 10^-4^). Bottom: Schematic of the subcellular location of condensates at the centrosome and at the cell periphery. **B.** Time lapse epifluorescence images of the delayed enrichment of mCh-5Fm in KIF1A condensates in a representative cell. The green and red arrows correspond to the nucleation of the condensate and the first visible enrichment of mCh-5Fm, respectively. Scale bar, 10 µm. **C.** Same as (B) for a KIFC1 condensate.

### Chemical induction of condensate transport and localization

Several endogenous condensates were found to interact and undergo transport along the cytoskeleton tracks^61^. Our motor-LLPS scaffolds could, by design, constitutively interact with microtubule fibers as soon as they are translated. We thus sought to examine the consequence of a sudden induction of the interaction between condensates randomly distributed through the cytosol and molecular motors. To this end, we devised an assay based on the rapamycin-dependent heterodimerization of FRB and FKBP (Fig. 4A). On one side we fused plus-end and minus-end motors to mCh-FRB (giving rise to KIF1A-mCh-FRB and BICD2-mCh-FRB, respectively) (Fig. 4A). On the other side, we fused our LLPS scaffold emGFP-5Fm to FKBP (FKBP-emGFP-5Fm) (Fig. 4A). We first analyzed the behavior of these proteins in the absence of rapamycin. After 24 h co-expression of FKBP-emGFP-5Fm and either KIF1A-mCh-FRB or BICD2-mCh-FRB, cells displayed several FKBP condensates, randomly dispersed in the cytosol and coexisting without interactions with FRB-fused motors (Fig. 4B). In some cells KIF1A-mCh-FRB accumulated at the cell periphery, while no particular enrichment of BICD2-mCh-FRB could be observed close to the nucleus (Fig. 4B).

**Figure 4:**
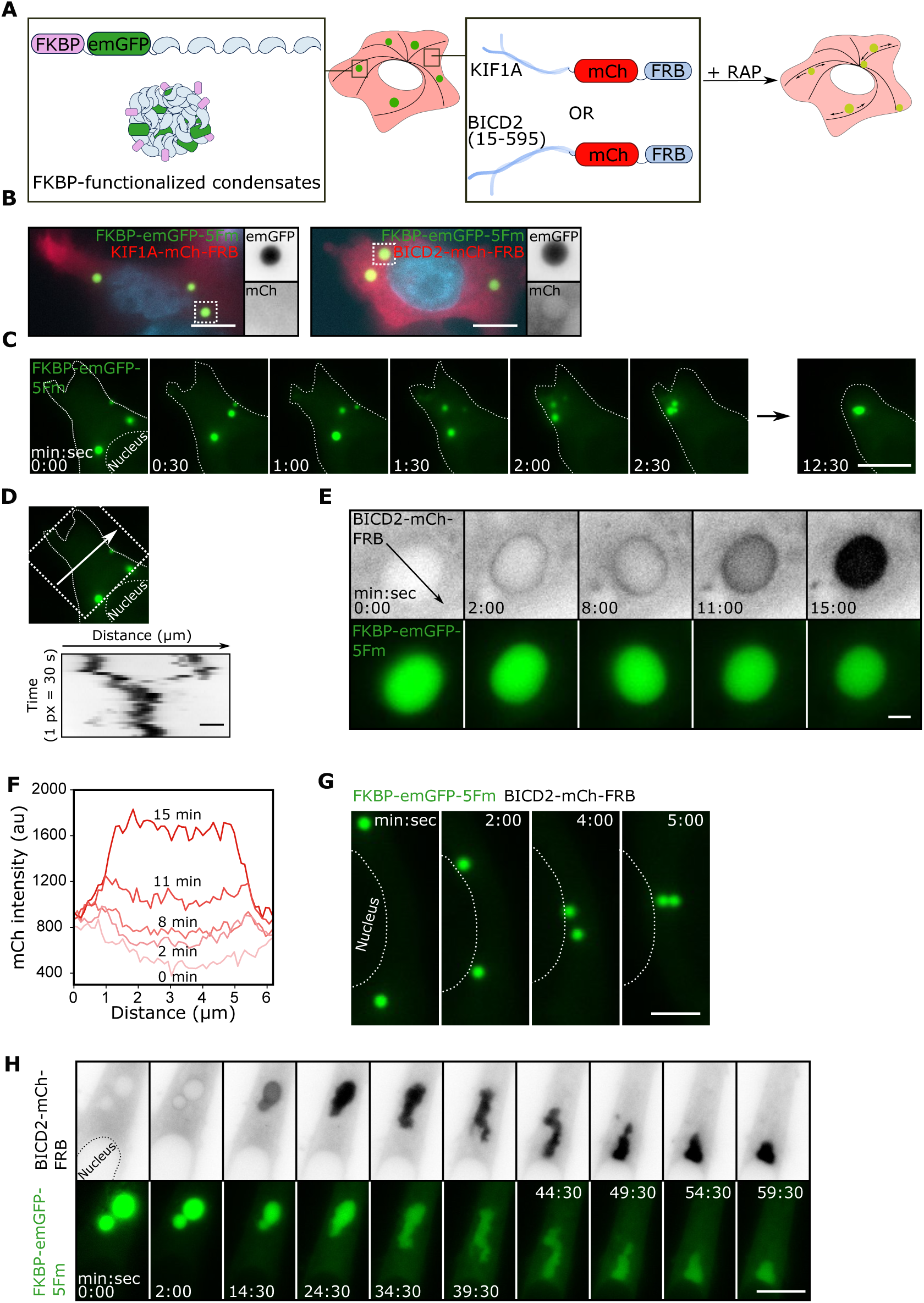
Chemically-induced binding of non-functionalized condensates to molecular motors led to their repositioning in cells. **A.** Schematic of the expected transport of FKBP condensates upon induction of their interaction with plus-end (KIF1A) or minus-end motors (dynein through BICD2) using Rapamycin (RAP) in HeLa cells. **B.** Representative epifluorescence imaging of cells expressing the FKBP-emGFP-5Fm LLPS scaffold and KIF1A-mCh-FRB (left) or BICD2-mCh-FRB (right) in the absence of rapamycin. Nuclei were stained with DAPI (blue). Grayscale inserts correspond to the green (emGFP) and red (mCh) channels of the region delineated by dashed squares. Scale bar, 10 µm. **C.** Time-lapse epifluorescence imaging of FKBP-emGFP-5Fm condensates undergoing transport towards the cell periphery and coalescence after addition of rapamycin at time 0. Scale bar, 10 µm. **D.** For the cell shown in (C), kymograph analysis along the 200 px-wide strip delineated by the arrows (∼ 21.7 µm), showing the coalescence of condensates over time. Scale bar, 2 µm. **E.** Epifluorescence imaging of the recruitment of BICD2-mCh-FRB (top) around a FKBP condensate (bottom), followed by progressive mixing of the two components. The black arrow corresponds to where the profile plots in (F) were plotted. Scale bar, 2 µm. **F.** Evolution of the mCherry intensity along the black arrow in **E** over time. **G.** Epifluorescence imaging of the transport of two FKBP condensates in a cell expressing BICD2-mCh-FRB after addition of rapamycin. Scale bar, 5 µm. **H.** Epifluorescence imaging of the recruitment and incorporation of BICD2-mCh-FRB (top) in a FKBP condensate (bottom) after addition of rapamycin, followed by transport towards the nuclear envelope with a liquid-like behavior. Scale bar, 10 µm.

We then added rapamycin (24 h after transfection) to induce interaction between the FKBP-condensates and KIF1A-mCh-FRB, and monitored the consequences using time-lapse microscopy. Within a couple of minutes, we could observe some events of long-range condensate transport toward the cell periphery. On Fig. 4C, we report an example of converging motions of condensates, which coalesced together at the cell extremity within a few minutes (Fig. 4D). The other cells, however, did not display obvious transport of condensates, which may be explained if initially the distribution of KIF1A motors was highly polarized towards the plasma membrane, making them unavailable for interaction with disperse condensates.

With BICD2-mCh-FRB, upon addition of rapamycin, we first observed the recruitment of the FRB-fused motor on the surface of the FKBP-emGFP-5Fm condensates, with a distinct mCherry corona forming in less than 1 min (Fig. 4E). Then, BICD2-mCh-FRB diffused towards the inner part of the FKBP condensates, driven by an internal mixing of the components, which occurred within 10 to 30 minutes (depending on the condensate size) (Fig. 4E and 4F). Subsequently, two types of directed motions towards the cell center were observed: some condensates were transported in a few minutes with no morphological change (Fig. 4G), while others underwent a striking deformation consistent with the rheological properties of a cytoplasm acting as a stiff and porous meshwork (Fig. 4H)^62^.

In conclusion, this assay allowed to chemically induce the rapid transport of condensates to either the cell periphery or the cell center.

### Localizing exogenous RNAs through the spatial positioning of condensates

The co-assembly of RNAs and RBPs into membrane-less organelles could potentially play a role in RNA trafficking to specific subcompartments or distal positions. Using a biomimetic approach, we thus sought to localize mRNAs by engineering motor condensates programmed to recruit a specific mRNA. Our strategy consisted of fusing MCP to our LLPS scaffold to enable the recruitment of RNAs with MS2 stem loops (Fig. 5A) ^40^. MCP scaffolds (MCP-5Fm) were then co-transfected with motor-LLPS scaffolds (KIF1A-emGFP-5Fm or BICD2-emGFP-5Fm), and with a plasmid expressing an RNA containing four MS2 repeats (RNA-MS2) (Fig. 5A).

**Figure 5:**
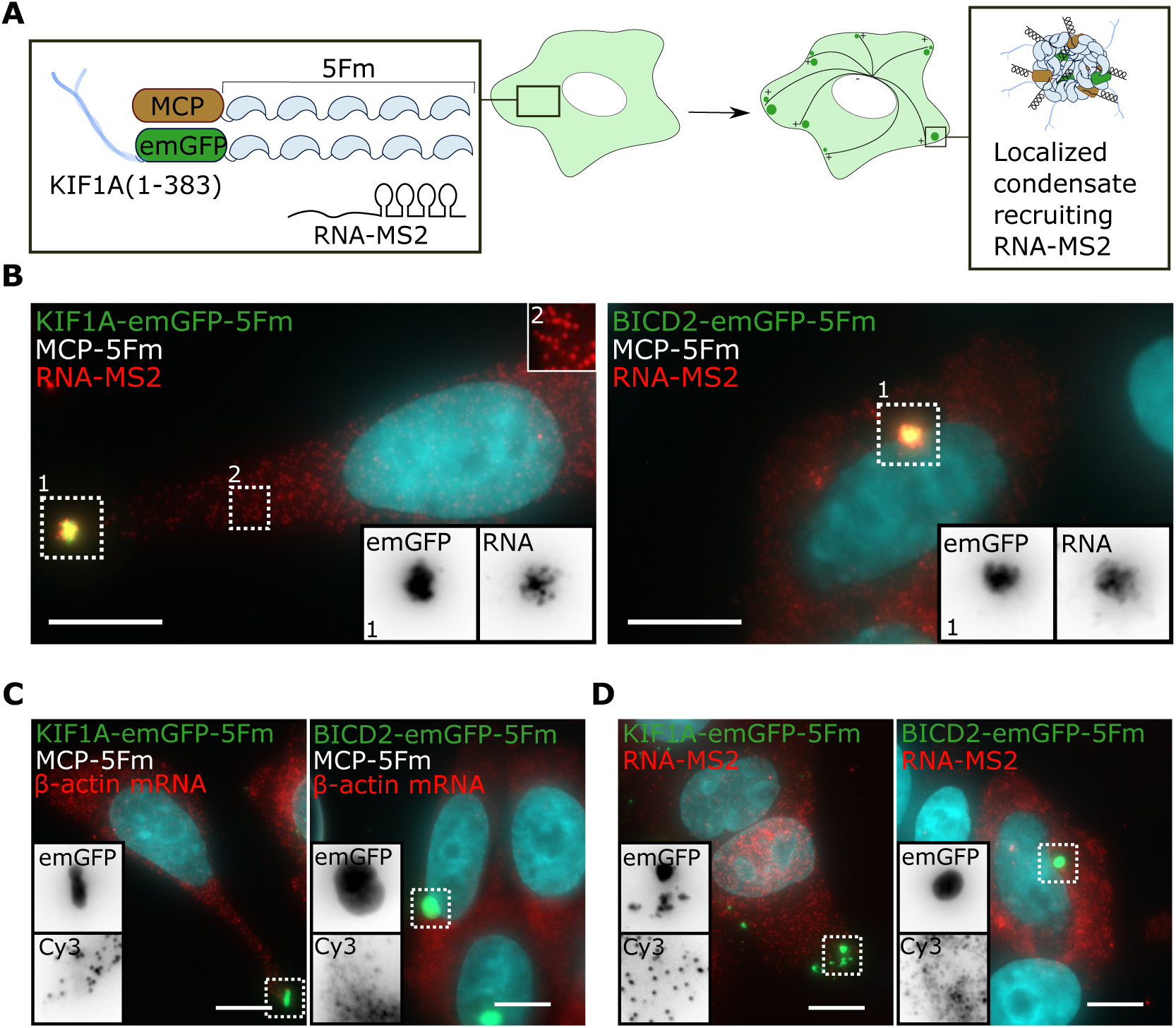
Motor/MCP condensates recruit RNA-MS2 and are efficiently positioned at the cell periphery or at the centrosome. **A.** Schematic of the formation of KIF1A/MCP condensates able to recruit the heterologous RNA-MS2. **B.** Representative epifluorescence imaging of cells containing KIF1A/MCP or BICD2/MCP condensates (green, left and right panels, respectively) following RNA-MS2 analysis by smiFISH (Cy3 probe, red). Nuclei were stained with DAPI (blue). The red channel setup allows for the visualization of dispersed RNA molecules while saturating the signal in the condensate. Grayscale inserts (1) correspond to the non-saturated green (emGFP) and red (Cy3) channels of the regions delineated by dashed squares. Insert 2 shows isolated RNA-MS2 molecules. Scale bar, 10 µm. **C.** Epifluorescence imaging of cells containing MCP condensates (green) following β-actin mRNA analysis by smiFISH (Cy3 probe, red). Scale bar, 10 µm. **D.** Epifluorescence imaging of cells containing condensates lacking MCP (green) following RNA-MS2 analysis by smiFISH (Cy3 probe, red). Scale bar, 10 µm.

We found that after 24 h of expression, bi-functionalized motor/MCP condensates were efficiently positioned at the cell periphery or at the centrosome depending on the motor’s directionality. Using single molecule FISH (smFISH), we demonstrated the recruitment of RNA-MS2 molecules in the motor condensates (each Cy3 dot corresponds to individual RNA molecule) (Fig. 5B). As specificity controls, the endogenous β-actin mRNA lacking MS2 stem loops was not recruited to MCP condensates (Fig. 5C), and the RNA-MS2 was not recruited on condensates lacking MCP (Fig. 5D). Therefore, condensates formed using a combination of motor- and MCP-LLPS scaffolds efficiently and specifically recruit MS2-containing RNAs.

Overall, these results demonstrate the specific localization of RNA via artificial condensates.

### Delocalizing endogenously tagged ASPM mRNA using motor condensates

To highlight a second application of our condensates, we aimed to use them as a tool to alter the subcellular localization of an endogenous RNA. To this end, we used a HeLa cell line in which 24 MS2 repeats were inserted in the 3’UTR of the Abnormal Spindle-like Microcephaly-associated (*ASPM*) gene using CRISPR-Cas9 (HeLa/ASPM-MS2). The resulting clone thus expresses the ASPM-MS2 mRNA in a stable manner, under the control of its endogenous promoter, which can be visualized by smFISH using a probe directed against the MS2 sequence^57^. Like untagged ASPM mRNA, ASPM-MS2 mRNA was weakly expressed during interphase and its expression increased during mitosis, with the mRNA localizing to centrosomes, particularly from early mitotic stages till metaphase (Fig. 6A)^57,63^. This created a striking local concentration of ASPM-MS2 mRNA on centrosomes making it an ideal candidate for delocalization attempts.

**Figure 6:**
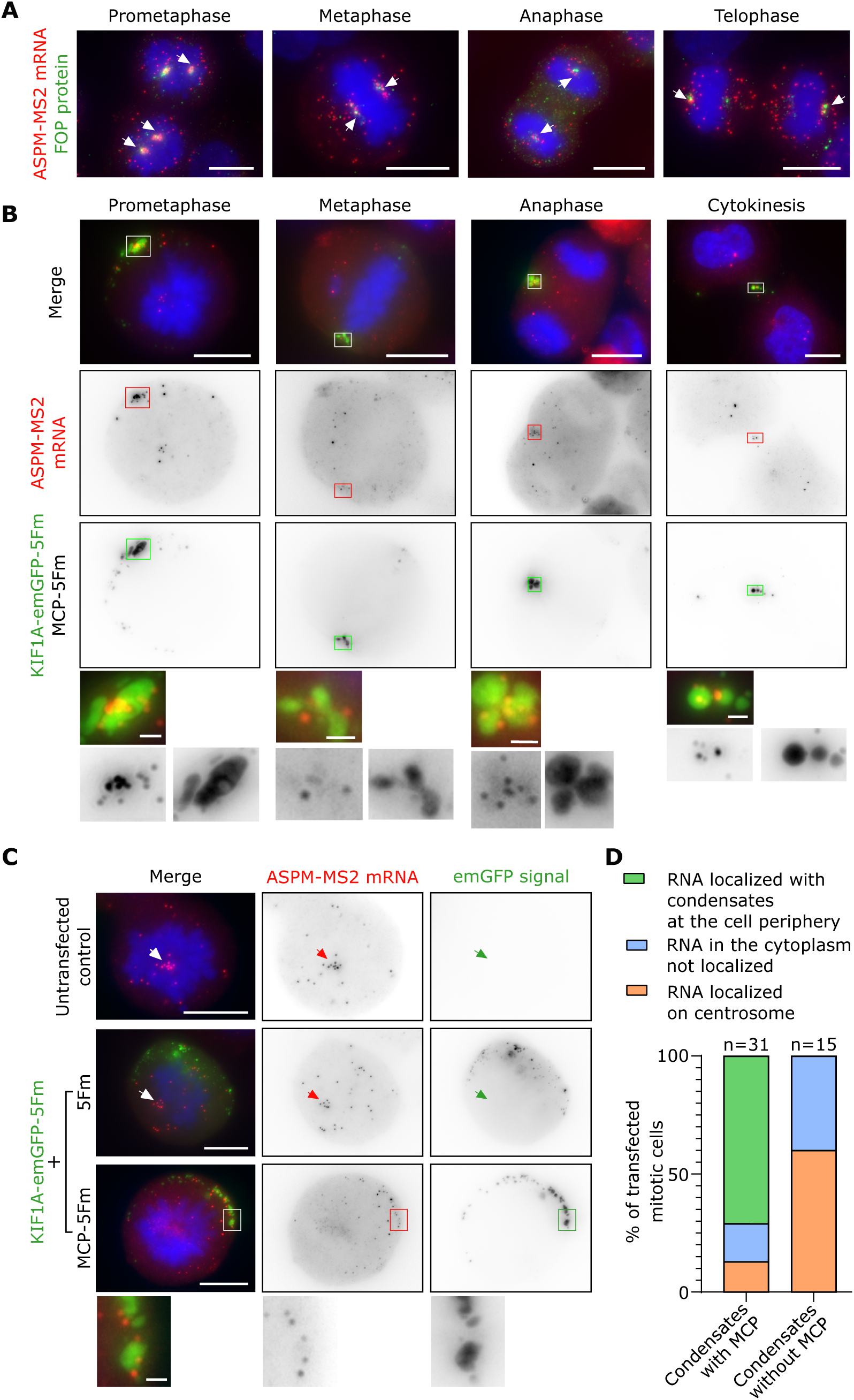
KIF1A condensates can efficiently delocalize ASPM-MS2 RNA towards the cell membrane during mitosis. **A.** Epifluorescence imaging of HeLa/ASPM-MS2 cells at different stages of mitosis after immunostaining of FOP (green) as a centrosome marker. The RNA was revealed by smFISH using a MS2 probe (red). DNA was stained with DAPI (blue). Scale bar, 10 µm. White arrows point to centrosomal mRNA accumulation. **B.** Epifluorescence imaging of HeLa/ASPM-MS2 containing KIF1A/MCP condensates at different stages of mitosis. Middle and bottom panels show the ASPM-MS2 mRNA revealed by smFISH (red channel) and the GFP condensates (green channel), respectively. Upper panels show the merged channels with DAPI-stained DNA in blue. Scale bar, 10 µm. Squares depicting areas near the cell membrane where granules and RNA co-localize are enlarged below. **C.** Epifluorescence imaging of prometaphase cells untransfected or expressing KIF1A condensates, with or without MCP. Left panels show the merged channels with DAPI-stained DNA in blue. Middle and right panels show the ASPM-MS2 mRNA revealed by smFISH (red channel) and the GFP condensates (green channel), respectively. Scale bar, 10 µm. Arrows point to centrosomal mRNA accumulation. Scale bar, 10 µm. The boxed area where the RNA and granules co-localize at the cell membrane, is enlarged below. Scale bar, 1 µm. **D.** For the two conditions shown in (C), bar graph representing the % of mitotic cells with ASPM-MS2 mRNA localized at the cell periphery, dispersed in the cytoplasm, or localized on centrosome (N= 31 and 15 cells, as indicated, each from two independent experiments).

To test this delocalization, we transiently transfected HeLa/ASPM-MS2 cells with our MCP and KIF1A scaffolds (MCP-5Fm with KIF1A-emGFP-5Fm). Remarkably, the KIF1A/MCP condensates successfully delocalized ASPM-MS2 mRNAs away from centrosomes towards the cell membrane across mitosis (Fig. 6B). As a negative control, we expressed condensates without MCP (5Fm only) functionalized with the KIF1A motor (KIF1A-emGFP-5Fm). In this condition, condensates localized at the cell periphery without recruiting ASPM-MS2 mRNA (Fig. 6C, D), thus confirming the specificity of the system. Moreover, we observed three patterns of KIF1A condensates in mitotic cells: local clustering of condensates at the membrane (i); or condensates distributed under the cell membrane producing either a half (ii) or a full (iii) crown pattern (Fig. S3). Interestingly, the ASPM-MS2 mRNA tended to distribute like the condensates, demonstrating the robustness of this tool (Fig. S3).

Conversely, we tested the possibility of forcing centrosomal localization of ASPM-MS2 mRNA in interphasic cells. First, as expected, condensates without motor (formed using emGFP-5Fm) were randomly localized in the cytoplasm and were able to recruit ASPM-MS2 mRNAs only in the presence of MCP-5Fm (Fig. S4A, B). In contrast, the BICD2 scaffold (BICD2-emGFP-5Fm) led to a single condensate at the centrosome, which was able to artificially localize some ASPM-MS2 mRNAs at the vicinity of centrosomes during interphase (Fig. S4C, D), at a time where the mRNA should not localize there.

Taken together, motor condensates are a versatile tool for altering the subcellular localization of RNA in living cells.

## DISCUSSION

How is the spatial positioning of biomolecular condensates orchestrated in cells? Whereas many mechanisms of spatial regulation have been described for membrane-bound organelles and other cargos, much less is known for condensates. Yet, despite the diversity of cytoplasmic RNP condensates, including RNA transport granules, stress granules, and P-bodies, one common feature relies on their interactions with microtubule-based cytoskeleton. In this study, we engineered artificial condensates functionalized with kinesin motor and dynein adaptor domains in order to examine their interplay with microtubules and its consequences on condensate formation and localization. We found that motor condensates were robustly positioned at the periphery of cells or at the vicinity of the centrosomes, as predicted from the direction of processivity of the motors. Next, we asked whether one could reconstitute a minimal RNP transport system to localize RNAs in cells. By incorporating MCP proteins into our motor condensates, we succeeded in recruiting MS2-tagged RNAs in asymmetrically positioned condensates.

In a first setting, LLPS scaffolds were directly fused to plus-end motors (KIF1A or KIF5B), or to minus-end motor / motor adaptor (KIFC1 or BICD2) and constitutively expressed in cells. Using this approach, we could investigate the formation of condensates made of proteins prone to phase-separate while interacting with microtubule fibers. Indeed, at early stage, motor-LLPS scaffolds underwent phase separation on microtubule fibers due to motor accumulation at the cell periphery or near the centrosome. One hypothesis is that the accumulation of motor-LLPS scaffolds on microtubule increases their local concentration which may account for their local condensation on microtubule surface lattice. The cooperative binding of the motor-LLPS scaffold on fibers, mediated by the repetitive nature of the LLPS scaffolds, could therefore favor prewetting on microtubules and phase separation below the expected saturation concentration ^64^. This process has recently been proposed for Tau and TPX2, two microtubule-associated proteins involved in the stabilization/nucleation of microtubule fibers^64^, or in a different context for the condensation of the transcription factor Klf4 on DNA molecules^65^.

To infer how nucleation of condensates was dependent on the capacity of the scaffolds to interact with microtubules, we monitored both motor-functionalized and non-functionalized LLPS scaffolds. Interestingly, we found that the condensation of the two LLPS scaffolds was sequential, with motor-LLPS scaffolds condensing systematically ahead of non-functionalized LLPS scaffolds. Non-functionalized scaffolds predominately accumulated at the sites of preformed motor condensates (Fig. 1D, 1F, 2E, 2F).

Classical nucleation theory predicts that phase-separated condensates can either form with no specific localization or, in contrast, at specific sites acting as seeds overcoming the kinetic barrier of nucleation. Recent studies showed how specific biomolecules can act as seeds and govern condensate nucleation at specific sites, such as DNA break sites^66^, the membrane^67–69^, or the apical side of the nucleus for nucleolus^70^. Our study provides an alternative scheme for the spatial positioning of nucleation. Here, we highlighted the positioning of condensates at polarity sites powered by microtubule-based motor proteins. In our system, condensate positioning occurred predominantly by nucleating phase separation at the destination sites of transported molecules rather than by transporting already formed condensates to their destination. This suggests a two-step mechanism (Fig. 7A): (i) active transport of the condensate scaffolds leading to their localization at polarity sites (microtubules extremities), (ii) nucleation of motor condensates through a mechanism possibly mediated by prewetting or cooperative binding. The pathway to such condensate localization is similar for the four motor domains studied. However, and in contrast to kinesin condensates, we also observed some events of nucleation of BICD2 condensates dispersed throughout the cytosol, which then were transported to the cell centrosome to eventually coalesce into a large condensate (Fig. 2F and S2C). This formation of BICD2 condensates in the cytosol prior to their transportation may be due to the requirement to assemble a high number of dyneins on the condensate surface to generate large collective forces and efficient transport^71^. Subsequently, recruitment of non-functionalized scaffolds in the preformed condensates could be observed.

**Figure 7:**
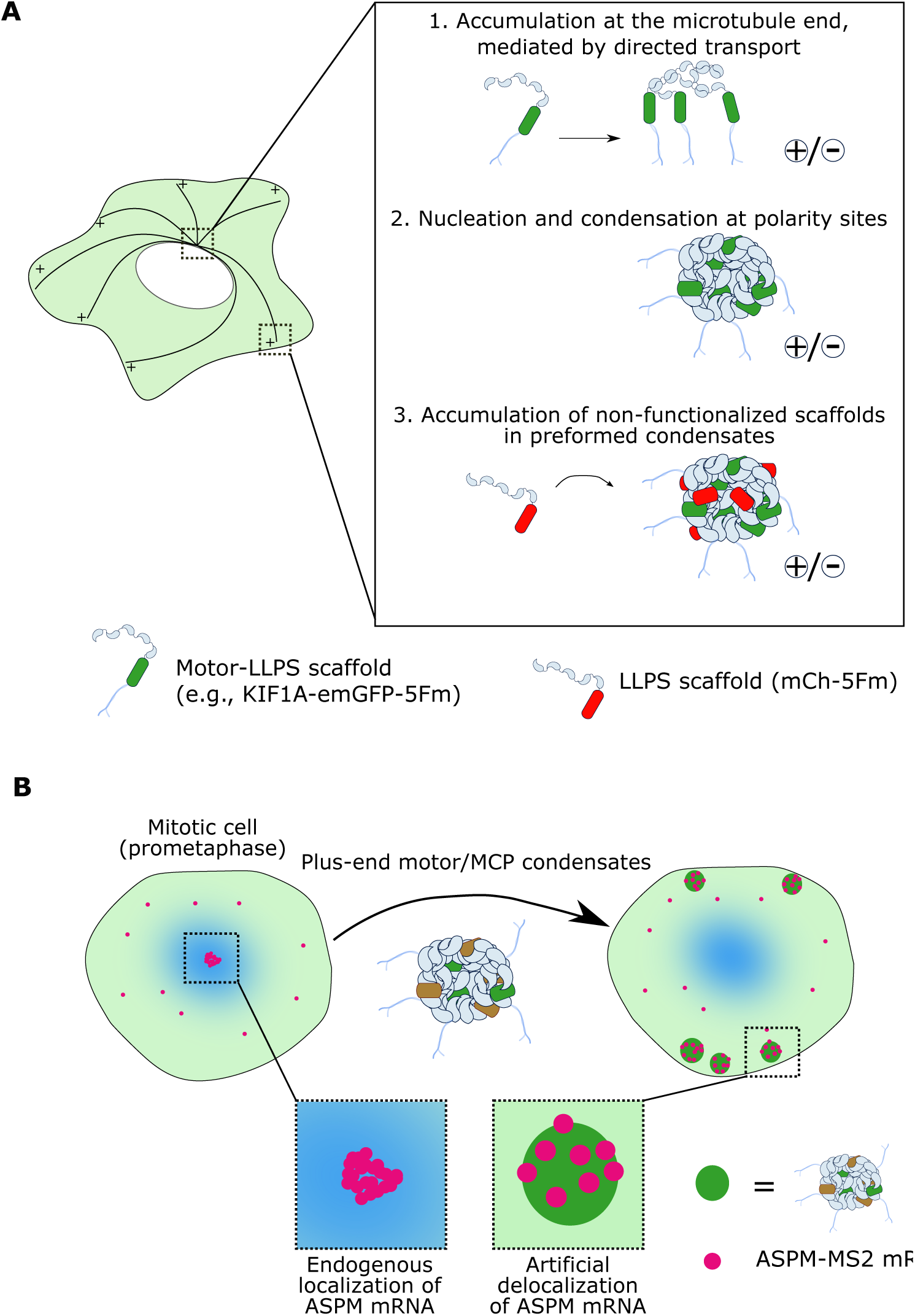
Model of localized nucleation and growth of condensates allowing for delocalization of ASPM-MS2 mRNA. **A.** Schematic model of the spatial localization of motor-functionalized condensates based on a stepwise mechanism: (1) active transport of the condensate scaffolds leading to their localization at microtubules extremities; (2) nucleation of motor condensates through a mechanism possibly mediated by prewetting or cooperative binding; (3) non-functionalized LLPS scaffolds accumulate in the preformed condensates. **B.** Artificial condensates drive the delocalization of individual ASPM-MS2 mRNAs at the cell periphery, suggesting that they outperform endogenous mRNA localization mechanisms by rewiring the transport machinery.

Coalescence of smaller BICD2 condensates into one larger condensate is reminiscent of the coalescence of stress granules upon transport along microtubules mediated by the dynein adaptor BICD1^6–8,10^. Previous studies on stress granules emphasized that their assembly follows distinct temporal steps, with first the formation of stable cores through multiple stable interactions, and secondly evolution into larger assemblies by recruiting a less dense shell^72^. Here, our studies highlight a simple mechanism based on LLPS where compositional complexity of granules builds during assembly processes in a sequential fashion. In our system, the localized nucleation of motor condensates provides a platform for the subsequent enrichment of non-functionalized LLPS scaffolds. This is reminiscent of the sequential localization of mRNAs observed during P-body formation in yeast^73^. Interestingly, in our system, the timing of enrichment of non-functionalized scaffolds into motor condensates depends on their localization in cells (Fig. 3). Therefore, site-specific nucleation combined with sequential enrichment provides a simple mechanism to build, in an ordered fashion, multicomponent condensates.

Other the last years, several chemical and optogenetic tools have been developed to perturb and control organelles positioning, interactions, and trafficking. Here, inspired by repositioning assays of membrane-bound organelles using chemically-induced dimerization strategy^74^, we extended our system to chemically trigger the interactions between dispersed condensates and microtubule motor proteins. With this approach, we obtained a temporal control of induction of condensate transport and localization at the cell periphery or at the vicinity of the centrosome (Fig. 4). Harnessing the trafficking of artificial condensates is a first step towards the assembly of biomimetic RNP transport system in cells.

Many RNAs are found localized in specific area of cells, and local translation is thought to participate to many functions dictating cell fate^13^. Complementary, mislocalization of RNA is reported to be associated with disease development^24^. There is consequently a strong emphasis to enlarge the current toolbox to analyze and study RNA localization and translation. The methodologies developed so far range from the visualization of RNP transport and translation with single molecule resolution, to spatial transcriptomics to map RNP interactomes^21,27,28,30,75,76^. In this context, we extended our assay to use it for the spatial manipulation of RNAs in cells. We showed that artificial condensates, functionalized with both motor domains and MCP proteins could be used as minimal RNP condensates recruiting a unique RNA, making it possible to explore condensate-mediated RNA delocalization. As a first proof-of-concept, we demonstrated the efficient recruitment of exogenous RNAs on motor condensates that were positioned at the cell periphery or at the centrosome depending on the motor directionality (Fig. 5). Combined with a temporal control of assembly/disassembly, one could anticipate future developments, where these artificial structures could act as platform organizing biochemistry in space and time.

Then, we demonstrated the ability of our system to strongly perturb the spatial distribution of endogenously tagged mRNAs. Artificial condensates drove the delocalization of individual ASPM-MS2 mRNAs at the cell periphery (Fig. 6,7B). This demonstrates how our system could outperform endogenous mRNA localization mechanisms by rewiring the transport machinery between the cytoplasm, the centrosome and the cell periphery. Competing with endogenous ASPM mRNA localization using artificial condensates provides interesting insights. On one hand, it has been shown that the ASPM RNA (as well as other centrosomal transcripts) naturally localizes to centrosomes through an active transport mechanism involving the microtubules and molecular motors^57^. This trafficking is dependent on the encoded nascent peptide and occurs rapidly at the onset of mitosis: within a couple of minutes, scattered RNA readily concentrates on centrosomes, as revealed by live imaging^57^. On the other hand, KIF1A condensates traffic away from the centrosomes and thus drag ASPM-MS2 mRNA. Since both the natural and artificial transport systems share microtubules for transit, the location where the RNA ends up provides an estimation of which localization process is more efficient. In most cells, the artificial condensates won the contest.

Several non-exclusive processes, that all rely on the capacity of KIF1A condensates to generate mechanical forces, could account for ASPM mRNA delocalization at the cell periphery: (i) Pulling forces applied by the condensates on individual ASPM mRNAs accumulated at the centrosome, allowing to convey RNAs along microtubule tracks. This suggests KIF1A forces are stronger than the cohesive forces bridging ASPM mRNA to centrosomal material. (ii) A tug-of-war between the KIF1A condensates and the endogenous transport machinery of ASPM mRNA to the centrosomes. For instance, KIF1A condensates can link individual RNAs to many more motors than a single nascent peptide or an endogenous adapter canonically would. They can be seen as a transport particle pulled by several molecular motors in a cooperative manner, allowing them to surpass the natural mechanism of ASPM mRNA transport. (iii) The direct capture and transportation of freely-diffusing ASMP RNAs by KIF1A condensates, upstream of their transport to the centrosome.

Interestingly, ASMP mRNA delocalization experiments provide a first benchmarking of the performance of our artificial condensates. This approach could open novel perspectives to examine the importance of RNA localization for cellular functions and may be extended to rewire the trafficking of other biomolecules of interest.

## MATERIAL AND METHODS

### Experimental model

Human epithelioid carcinoma HeLa cells (ATCC, ccl-2) were kept in Dulbecco’s modified Eagle’s medium (with 4.5 g/L D-glucose, HyClone) with 10% fetal bovine serum (Gibco, 10,270,106) and 1% Penicillin-Streptomycin (Sigma, P4333), at 37°C in a 5% CO_2_ humidified atmosphere. Tests for mycoplasma contamination were routinely carried out.

### Plasmids

To generate the constructs containing 5 repeats of FKPB-F36M, a first plasmid puCIDT-Amp-5Fm was designed containing five repeats of FKBP-F36M separated by sequences coding for linkers of four GGS repeats (12 amino acids total). To avoid recombination, degenerate repeats were used. The first repeat was preceded by a Nhe and an AfeI restriction sites and the last one was followed by a Xba1 restriction site. This plasmid was purchased from IDT. To obtain the pcDNA3.1-5Fm plasmid (called hereafter 5Fm), puCIDT-Amp-5Fm was digested with NheI and XbaI, and the 5Fm containing fragment was subcloned between NheI and XbaI sites in the pcDNA3.1 (+) vector (Invitrogen). pcDNA3.1-emGFP-5Fm, pcDNA3.1-mCh-5Fm (called hereafter mCh-5Fm) and pcDNA3.1-MCP-5Fm (called hereafter MCP-5Fm) were then obtained by inserting emGFP, mCherry or a tandem MCP coding sequence, respectively, between HindIII and AfeI restriction sites.

Coding sequences for human KIF1A(1-383), mouse KIF5B(1-555), human KIFC1(125-673) and mouse BICD2(15-595) were obtained from Addgene (plasmids #133242, #120170 #120169, and #120168 respectively)^60,77^. KIF1A and KIF5B coding sequences were inserted in pcDNA3.1-5Fm between NheI and AfeI restriction sites, with respectively EcoR1 and Not1 restriction sites ahead of AfeI for subsequent sub-cloning. Then emGFP was inserted in pcDNA3.1-KIF1A-5Fm between EcoRI and AfeI, and in pcDNA3.1-KIF5B-5Fm between NotI and AfeI restriction sites. KIFC1 coding sequence was inserted in pcDNA3.1-emGFP-5Fm between XbaI and AgeI restriction sites. pcDNA3.1-BICD2-emGFP-5Fm plasmid was obtained by adding a NheI restriction site ahead of emGFP in pcDNA3.1-emGFP-5Fm, and inserting the BICD2 coding sequence between HindIII and NheI restriction sites. The four pcDNA3.1 constructs KIF1A-emGFP-5Fm, KIF5B-emGFP-5Fm, emGFP-KIFC1-5Fm and BICD2-emGFP-5Fm, are hereafter called the motor-LLPS scaffolds.

### Transfection

For imaging after cell fixation, HeLa cells were cultured on 22×22 mm glass coverslips (VWR) in 6-well plates (Falcon, 3.5×10^5^ cells per well). For live imaging, HeLa cells were seeded on 35-mm-dishes with polymer coverslip bottom (Ibidi, 1.5×10^5^ cells per dish). For both, cells were transfected 24 hours later using Lipofectamine 2000 (Invitrogen) according to the manufacturer’s protocol. For fixed cell imaging, cells were transfected with a 2:1:1 ratio of 5Fm, mCh-5Fm and motor-LLPS scaffold (2 µg total per well). The same conditions were followed for control experiments with motors lacking the LLPS 5Fm domain. In the case of KIFC1, cells were transfected with a modified ratio of 2.5:1:0.5. For live imaging (formation, dissolution and induction acquisitions), cells were transfected with a 1:1 ratio of motor-LLPS and 5Fm scaffolds (800 ng total per µ-dish). For smFISH experiments, cells were transfected with a 1:1:2 ratio of motor-5Fm scaffold, 5Fm, and MCP-5Fm (2 µg total per well) and 50 ng of RNA-MS2 plasmid. The ratio was modified to 0.5:1.5:2 in the case of KIFC1.

To probe condensate inhibition and dissolution (Fig. S1 and S2), FK506 (Sigma, F4679) was used at 2.5 µM. For chemical induction experiments of condensate transport (Fig. 3), rapamycin was used at 0.4 µM.

### ASPM-MS2 cell line generation

HeLa Kyoto cells were transfected with a combination of plasmids expressing the Cas9-nickase protein, two guide RNAs targeting the end of the ASPM gene, and a repair template harbouring 500 nucleotide homology arms. Homology arms flanked 3 HA tags, a stop codon, 24 MS2 repeats and an IRES-NeoR-stop codon sequence. This repair template was designed to allow insertion at the endogenous ASPM stop codon. Following neomycin selection at 400 μg/ml for 7-10 days, clones were isolated and characterized by PCR genotyping and smFISH to ensure proper cassette insertion and edited RNA localization. The clone used in this study is heterozygous as described in detail in Safieddine et al^57^. The sequences targeted by the guide RNAs are: TCTCTTCTCAAAACCCAATCtgg for guide 1, and GCAAGCTATTCAAATGGTGAtgg for guide 2, where lowercase corresponds to PAM sequences.

### Single-molecule fluorescence in situ hybridization

Single RNA molecule detection of the heterelogously expressed RNA-MS2 was performed according to the previously described smiFISH (single-molecule inexpensive FISH) method^78^. Briefly, cells were fixed in 4% paraformaldehyde (PFA) for 20 min at RT, and permeabilized with 70% ethanol in phosphate buffer saline (PBS) at 4°C overnight. A mix of gene specific (described previously^54^) and Cy3 FLAP probes in hybridization buffer (50 µl/coverslip) was used for overnight hybridization at 37°C in a humidity chamber. After washing twice for 30 min at 37°C in 15% formamide in SSC buffer and rinsing twice in PBS, cells were either mounted with VECTASHIELD mounting medium containing DAPI (Vector Laboratories, H-1200) or processed through immunofluorescence steps.

smFISH against the MS2 sequence in HeLa/ASPM-MS2 cells was done using a single probe (25 ng of probe per 100 μl of hybridization mixture) that had the following sequence: 5’AT*GTCGACCTGCAGACAT*GGGTGATCCTCAT*GTTTTCTAGGCAATT*A where * denotes a thymidine conjugated with a Cy3 molecule. Hybridization was done on cells grown on a glass coverslip in a buffer containing 20% formamide (Sigma-Aldrich), 1x SSC, 0.34 mg/ml tRNA (Sigma-Aldrich), 2 mM VRC (Sigma-Aldrich), 0.2 mg/ml RNAse-free bovine serum albumin (BSA, Roche Diagnostics) and 10% dextran sulfate (MP Biomedicals). Hybridization was done overnight at 37°C and coverslips were washed the next day in a 20% formamide 1x SSC solution twice, each at least for 40 mins at 37°C. Coverslips were then mounted using VECTASHIELD containing DAPI (Vector Laboratories, H-1200).

### Immunofluorescence

For centrosome imaging in Figure 2, cells were fixed 24 h after transfection in methanol at - 20°C for 10 minutes. They were then permeabilized with a solution of 0.2% Triton X-100 and 0.1% BSA in PBS for 30 min, incubated for 1 h with the primary antibody (rabbit anti-pericentrin, Covance PRB-432C, 1:500 dilution), washed three times with PBS at RT for 5 min, incubated for 1 h with the secondary antibody (AffiniPure Goat Anti-Rabbit conjugated with Alexa Fluor 488 dye), washed three times with PBS at RT for 5 min, and finally mounted with VECTASHIELD containing DAPI (Vector Laboratories, H-1200).

### Imaging

For live imaging, cells were imaged on a Zeiss LSM 710 META laser scanning confocal microscope using an x63 oil-immersion objective (PlanApochromatic, numerical aperture (NA) 1.4), at 37°C in a 5% CO2 humidified atmosphere, starting 4 h after transfection (formation). Microscope hardware and image acquisition were controlled with LSM Software Zen 2012. For fixed experiments, cells were imaged using an IX81 microscope (Olympus) and 60x oil immersion objective (PlanApo, NA 1.42), equipped with a CMOS camera, Orca-Fusion (Hamamatsu, Corporation), and a LED system of illumination (Spectra X, Lumencor). Microscope settings were controlled using Micro-manager on ImageJ. Images were analyzed using Fiji.

### Data analysis

To quantify the degree of asymmetry of the condensate distribution, the fraction of mCh-5Fm fluorescence in the peripheral 25% of the cells (I_25_) was measured by adapting a previously published method^58^. For each cell, a first circle encompassing the entire cell and having for center a point in the nucleus was drawn. A concentric circle with a 10-pixel diameter was then drawn, from which a series of concentric circles were derived by iteratively enlarging the diameter by 10 pixels until reaching the first circle size. 5-pixel rings were then built by subtracting each circle to the next one in the series. Final regions of interest (ROI) were defined as the overlap between each ring and the cytoplasm. The images were subsequently processed by applying Gaussian blur (5-pixel radius) to eliminate strong local variations in intensity. For each ROI, the area and the mean and minimal intensities were measured. Minimal intensities were subtracted to mean values to remove background, and integrated intensity were calculated for each ROI. The peripheral 25% of a cell was defined as the sum of consecutive ROI, starting from the most peripheral one, reaching 25% of the total cell area. For each cell, I_25_ was obtained by calculating the ratio between the integrated fluorescence of the mCherry signal in the peripheral 25% over the integrated fluorescence of the entire cell.

### Statistical analysis

For Fig. 1C and 3A, Wilcoxon rank-sum tests (nonparametric test to compare two distributions) were performed using the ranksum MATLAB function (MathWorks). For Fig. 2C, Pearson’s chi-squared tests (nonparametric test for nominal variables) were performed using the Python’s chi2_contingency function.

## Acknowledgments

We thank M.A. Plamont for her help along the project and C. Amari for his help in the implementation of the quantification process. We thank A. Meunier for providing the pericentrin antibodies and E. Bertrand for sharing HeLa/ASPM-MS2 cells.

## Funding

A.C was supported by an IPV-SU PhD fellowship. A. S. was supported by ANR (ANR-19-CE12-0024-01). P.C was supported by a FRM (MND202003011470) grant. This work was supported by the CNRS, Ecole Normale Supérieure, and ARC (20181208003) to Z.G, FRM (MND202003011470) and iBio (SU) to Z.G and D.W, and ANR (ANR-19-CE12-0024-01) to D.W.

## Author Contributions

A.C and Z.G conceived the project. A.C and P.C designed the motor condensate toolbox, as well as performed and analyzed motor-condensate experiments. A.S, M-N.B and D.W performed and analyzed ASPM-related experiments. A.C and Z.G prepared the manuscript with contributions from A.S, P.C and D.W. All authors discussed the results and commented on the manuscript.

## Declaration of Interests

The authors declare that they have no competing interests.

## Supporting Information

4 Supplementary Figures and 4 Supplementary Movies.

---

**Figure S1:**
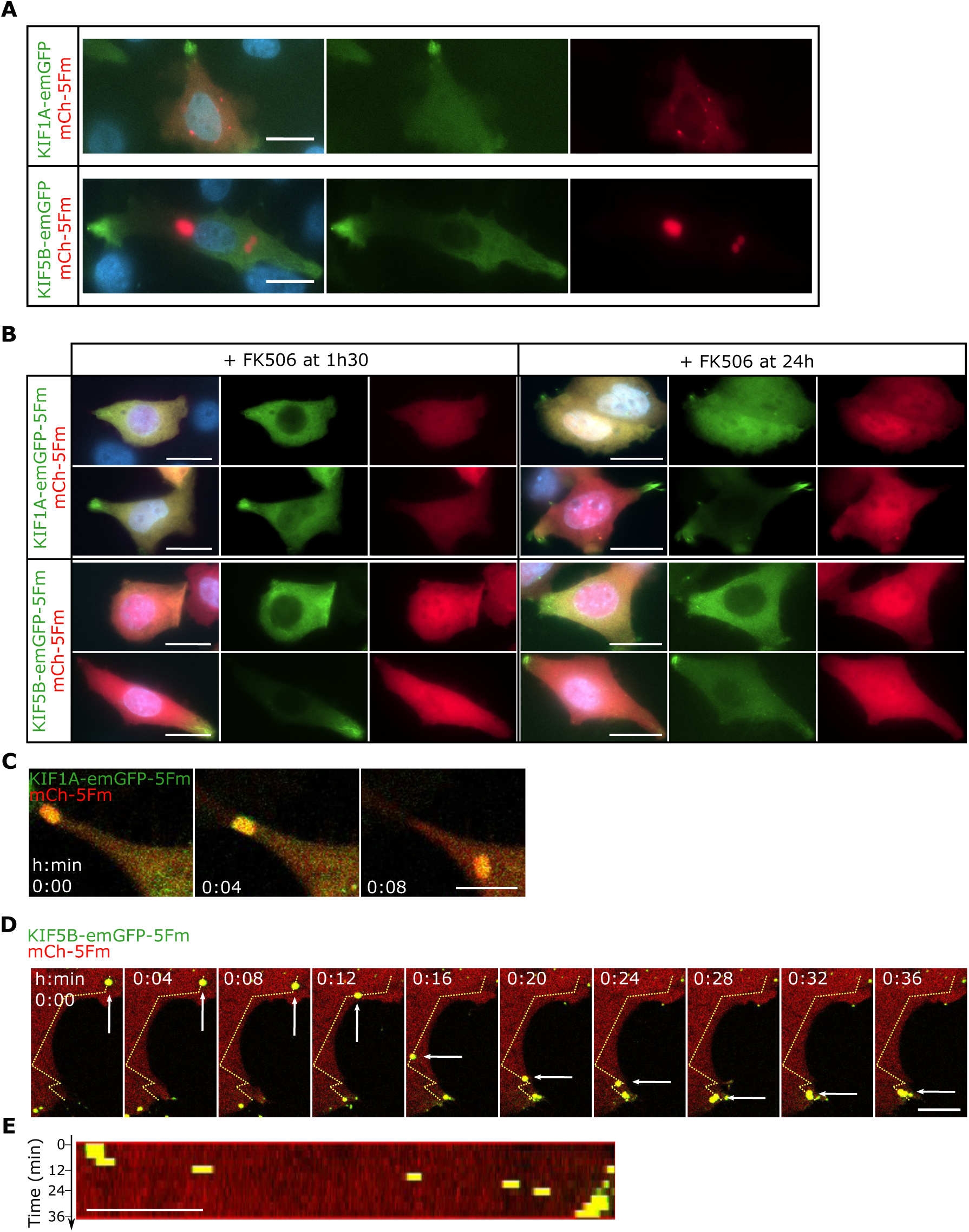
Cells expressing plus-end motors lacking the LLPS domain; chemical inhibition of condensates; and directed transport of condensates. **A.** Representative epifluorescence imaging of HeLa cells transfected with KIF1A-emGFP or KIF5B-emGFP (without LLPS domain) and mCh-5Fm. Nuclei were stained with DAPI (blue in merge). Scale bar, 20 µm. **B.** Epifluorescence imaging of HeLa cells expressing KIF1A- or KIF5B-emGFP-5Fm and mCh-5Fm after FK506 addition either right after transfection to forestall the formation of condensates (left) or 24 h after transfection to dissolve the condensates (right) (two examples for each condition). Nuclei were stained with DAPI (blue in merge). Scale bar, 20 µm. **C.** Directed transport of a KIF1A condensate. Scale bar, 10 µm. **D.** Directed transport of a KIF5B condensate (white arrows). More time points are given below for the area delimited by the white dashed rectangle. The dashed yellow line represents the condensate trajectory. Scale bar, 10 µm. **E.** Kymograph analysis along the condensate trajectory shown in (C). Scale bar, 10 µm.

**Figure S2:**
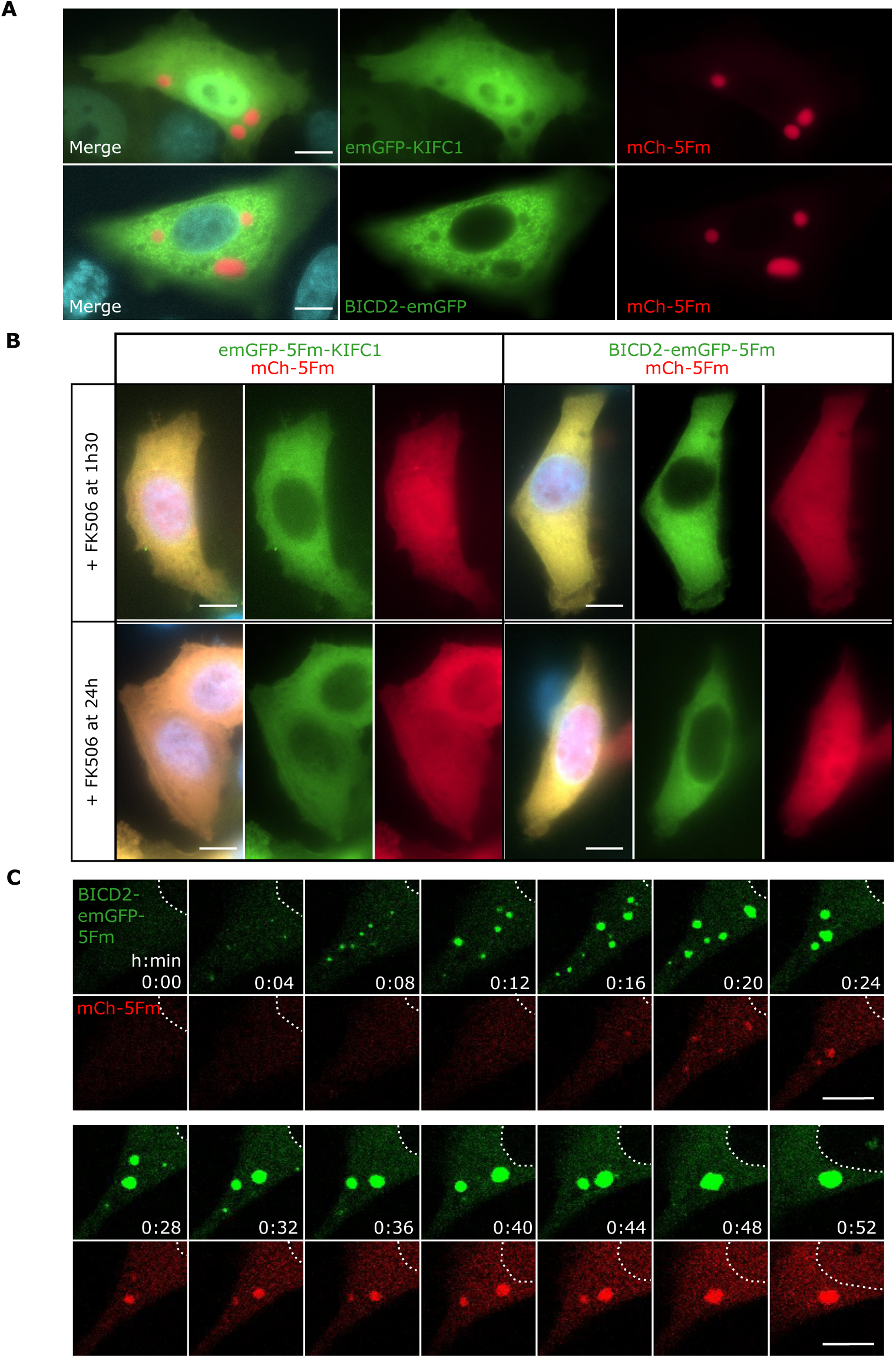
Cells expressing minus-end motors lacking the LLPS domain; chemical inhibition of condensates; and coalescence of BICD2 condensates. **A.** Representative epifluorescence imaging of HeLa cells transfected with emGFP-KIFC1 (upper panel) or BICD2-emGFP (lower panel) and mCh-5Fm. Nuclei were stained with DAPI (blue in merge). Scale bar, 10 µm. **B.** Epifluorescence imaging of HeLa cells expressing emGFP-5Fm-KIFC1 (left) or BICD2-emGFP-5Fm (right) and mCh-5Fm after FK506 addition either right after transfection to forestall the formation of condensates (top) or 24 h after transfection to dissolve the condensates (bottom). Nuclei were stained with DAPI (blue in merge). Scale bar, 10 µm. **C.** Time-lapse confocal imaging of occasionally dispersed nucleation of BICD2 condensates, followed by a delayed enrichment of the non-functionalized LLPS scaffold, directed transport towards the centrosome and coalescence. The white dashed line delineates the nucleus. Scale bar, 10 µm.

**Figure S3:**
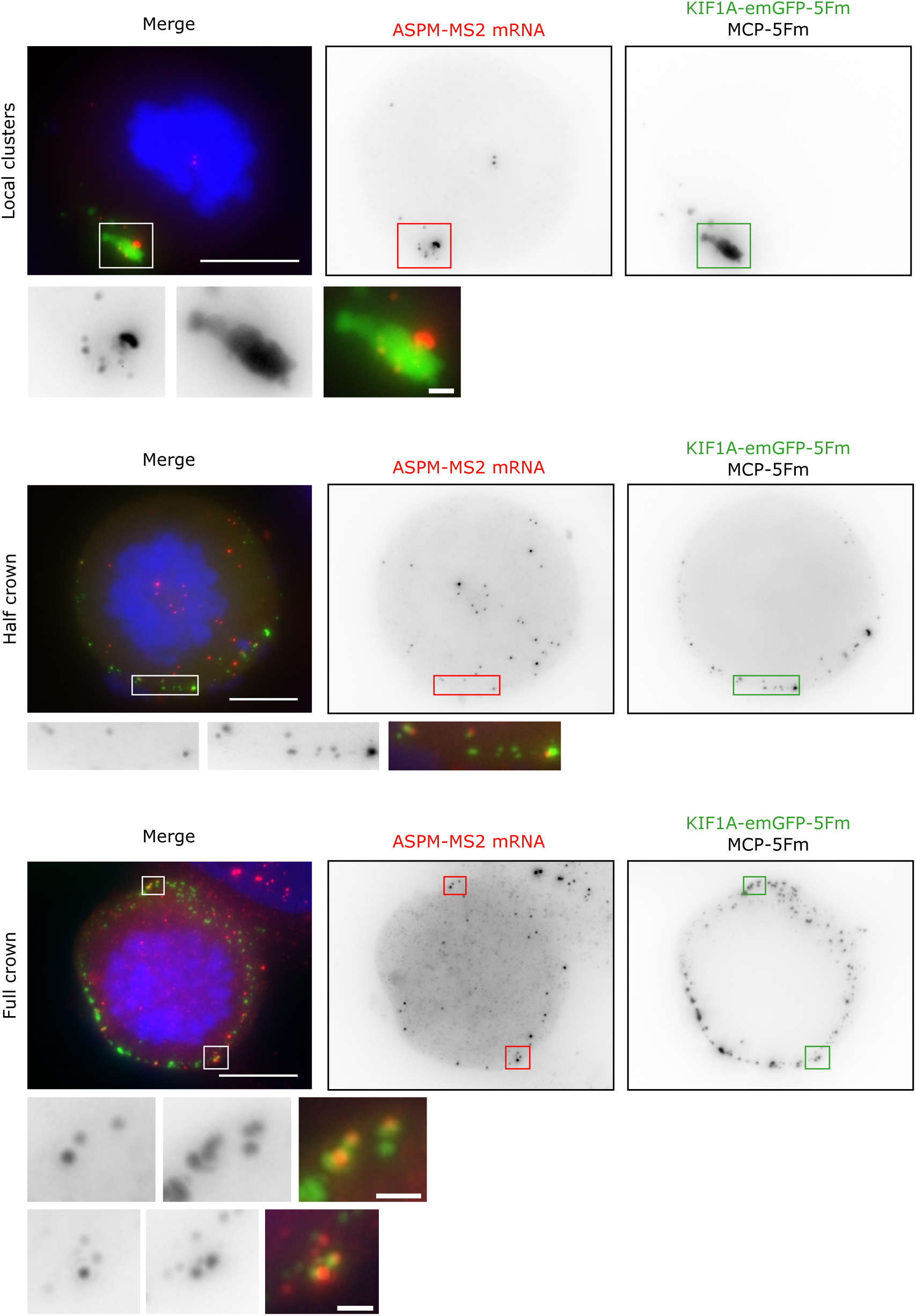
KIF1A/MCP condensates delocalized ASPM-MS2 mRNAs away from centrosomes and display three patterns in mitotic cells. Epifluorescence imaging of prometaphasic HeLa/ASPM-MS2 cells containing KIF1A/MCP condensates. Left panels show the merged channels with DAPI-stained DNA in blue. Left and middle panels show the ASPM-MS2 mRNA revealed by smFISH (red channel) and the GFP condensates (green channel), respectively. Scale bar, 10 µm. Squares depicting areas near the cell membrane where granules and RNA co-localize are enlarged below. Scale bar, 1 µm.

**Figure S4:**
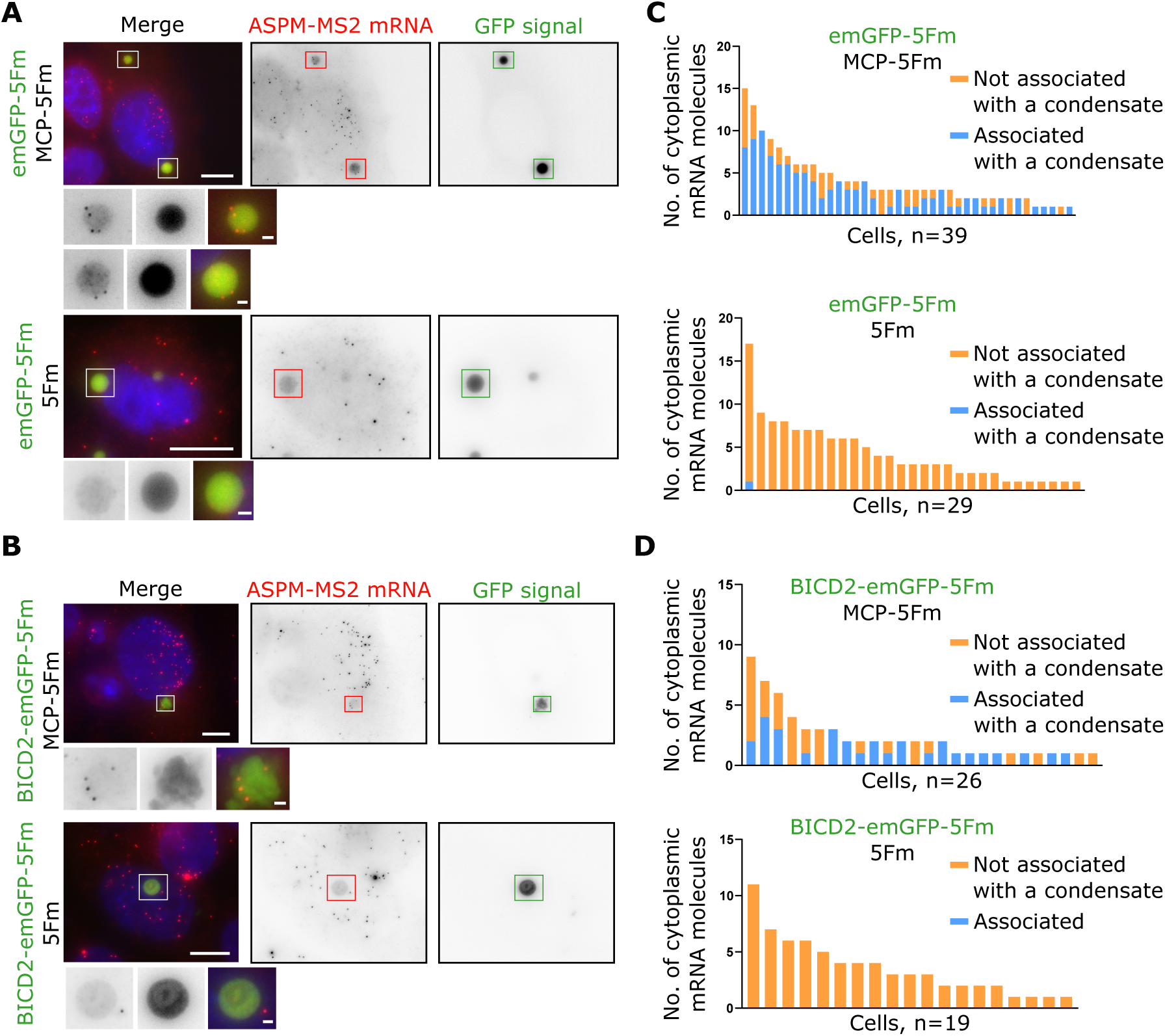
Motor-free BICD2 condensates can recruit ASPM-MS2 RNA in interphase. **A.** Epifluorescence imaging of interphasic HeLa/ASPM-MS2 cells containing motor-free condensates. Left and middle panels show the ASPM-MS2 mRNA revealed by smFISH (red channel) and the GFP condensates (green channel), respectively, either with (upper panels) or without (lower panels) MCP. Right panels show the merged channels with DAPI-stained nuclei in blue. Scale bar, 10 µm. Squares containing condensates that may (upper panels) or may not (lower panels) contain RNA are enlarged below. Scale bar, 1 µm. **B.** For the two conditions shown in (A), the bar graph shows the number of ASPM-MS2 RNAs per cell co-localizing or not with a condensate (N=39 and 29 cells, as indicated, each from two independent experiments). **C, D.** Same as in (A, B) for condensates containing the BICD2 motor adaptor.

## Notes

### Competing Interest Statement

The authors have declared no competing interest.

